# Effects of daily L-dopa administration on learning and brain structure in older adults undergoing cognitive training: a randomised clinical trial

**DOI:** 10.1101/482679

**Authors:** Alexander V. Lebedev, Jonna Nilsson, Joanna Lindström, William Fredborg, Ulrika Akenine, Carolina Hillilä, Pia Andersen, Gabriela Spulber, Elizabeth CM de Lange, Dirk-Jan van den Berg, Miia Kivipelto, Martin Lövdén

**Affiliations:** Aging Research Center, Department of Neurobiology, Care sciences, and Society, Karolinska Institutet, Stockholm, Sweden (A.V. Lebedev MD PhD, J. Nilsson PhD, Prof. M. Kivipelto MD PhD, Prof. M. Lövdén PhD, J. Lindström, W. Fredborg); Department of Clinical Neuroscience (CNS), Karolinska Institutet, Stockholm, Sweden (A.V. Lebedev MD PhD); Division of Clinical Geriatrics, Department of Neurobiology, Care Sciences and Society (NVS), Karolinska Institutet, Stockholm, Sweden (Prof. M. Kivipelto MD PhD, U. Akenine, C. Hillilä, P. Andersen MD, G. Spulber MD PhD); Theme Aging, Karolinska University Hospital, Stockholm, Sweden (Prof. M. Kivipelto MD PhD, U. Akenine, C. Hillilä, P. Andersen MD, G. Spulber MD PhD); Neuroepidemiology and Ageing Research Unit, School of Public Health, Imperial College London, London, United Kingdom (Prof. M. Kivipelto MD PhD); Department of Neurology, Institute of Clinical Medicine and Institute of Public Health and Clinical Nutrition, University of Eastern Finland, Kuopio, Finland (Prof. M. Kivipelto MD PhD); Leiden Academic Centre for Drug Research, Division of Systems Biomedicine and Pharmacology, Universiteit Leiden, Leiden, Netherlands (Prof. E.C.M. de Lange PhD, D-J. van den Berg)

## Abstract

Cognitive aging creates major individual and societal burden, motivating search for treatment and preventive care strategies. Behavioural interventions can improve cognitive performance in older age, but effects are small. Basic research has implicated dopaminergic signaling in plasticity. We investigated whether transient enhancement of dopaminergic neurotransmission via administration of L-dopa improves effects of cognitive training on performance.

Sixty-three participants for this randomised, parallel-group, double-blind, placebo-controlled trial were recruited via newspaper advertisements. Inclusion criteria were: age of 65-75 years, Mini-Mental State Examination score >25, absence of serious medical conditions. Eligible subjects were randomly allocated to either receive 100/25mg L-dopa/benserazide (*n*=32) or placebo (*n*=31) prior to each of twenty cognitive training sessions administered during a four-week period. Participants and staff were blinded to group assignment. Primary outcomes were latent variables of spatial and verbal fluid intelligence.

Compared to the placebo group, subjects receiving L-dopa improved less in spatial intelligence (−0.267 SDs; 95%CI [−0.498, −0.036]; p=0.024). Change in verbal intelligence did not significantly differ between the groups (−0.081 SDs, 95%CI [−0.242, 0.080]; p=0.323). Subjects receiving L-dopa also progressed slower through the training and the groups displayed differential volumetric changes in the midbrain. Adverse events occurred for 10 (31%) and 7 (23%) participants in the active and control groups, correspondingly. No statistically significant differences were found for the secondary cognitive outcomes.

The results speak against early pharmacological interventions in older healthy adults to improve broader cognitive functions by targeting the dopaminergic system and provide no support for learning-enhancing properties of L-dopa supplements. The findings warrant closer investigation about the cognitive effects of early dopamine-replacement therapy in neurological disorders. This trial was preregistered at the European Clinical Trial Registry, EudraCT#2016-000891-54.

**Significance:** The results put constraints on the hypothesis of a key role of the deteriorated dopaminergic system in age-related decline of learning abilities, and speak against early pharmacological interventions in older healthy adults to improve cognitive functions by targeting the dopaminergic system. Our findings also raise concerns about usefulness of novel L-dopa-containing supplements that claim to have neuroprotective and learning-enhancing properties, and present an urgent need to carefully investigate the cognitive outcomes of early pro-dopaminergic interventions in clinical populations often receiving substantially larger doses of L-dopa.

## INTRODUCTION

Age-related cognitive decline and dementia are serious public health problems with devastating impact on the quality of life of individuals, their caregivers, but also on healthcare in general. The current total worldwide cost of dementia is about a trillion US dollars a year and is expected to double by 2030 (1). Whilst pharmacological treatment approaches have been unsuccessful, recent studies have demonstrated that combining cognitive training with exercise and a healthy diet can affect cognitive functioning in at-risk older people (2). Both exercise and dietary nutrients have been put forward as enhancers of neurobiological plasticity (3, 4), and may therefore increase effectiveness of cognitive training. One of the mechanisms through which exercise and diet are thought to modulate brain plasticity is through their action on dopaminergic neurotransmission (5, 6). Indeed, several independent lines of basic research have demonstrated the involvement of dopamine signaling in learning (7–11) and neurobiological plasticity (12, 13). Furthermore, plasticity has been shown to be reduced in healthy aging (14) and linked to changes in dopamine levels (15). Importantly, temporary augmentation of diminished dopaminergic neurotransmission can be accomplished by administering the catecholamine precursor L-dopa with beneficial effects on cognitive performance and learning in healthy adults (7, 8, 16–18). We aimed to bridge the knowledge gap between clinical and basic sciences by isolating and studying the role of dopamine system in learning and neurobiological plasticity in a sample of older adults drawn from the general population. In order to systematically implement this in the experimental setting, we designed the present randomised placebo-controlled clinical trial centred on transfer effects of working memory training to fluid intelligence (19, 20), which was selected as a primary outcome that may pick up task-independent training effects on working memory ability due to the important role that this ability plays in solving fluid intelligence task (21). Specifically, we hypothesised that enhancing dopaminergic neurotransmission, transiently during cognitive training, by administering L-dopa would lead to increased efficiency of cognitive training for improving general cognitive performance in older age.

## METHODS

### Study design and participants

For the present randomised, parallel-group, double-blind, placebo-controlled trial conducted at the Karolinska University Hospital in Huddinge, Stockholm, Sweden, healthy older individuals aged 65–75 years were recruited via daily newspaper advertisement. Eligibility criteria were initially assessed via a telephone screening and later during introduction meetings. Inclusion criteria were a Mini-Mental State Examination score of at least 26 points, absence of any serious medical or psychiatric conditions, no history of brain injuries or serious head traumas, no metal implants hindering Magnetic Resonance Imaging (MRI), no previous participation in studies employing cognitive training, right-handedness, and absence of colour-blindness. The study protocol (see “Main documents” at https://osf.io/nwwx8/) was approved by the regional ethics review board in Stockholm (2016-1897-31/1) and the Swedish medical product agency (20016-000891-54). Participants provided written informed consent before enrolment.

### Randomisation and Blinding

After a medical screening and baseline cognitive assessments conducted by experienced physicians, research assistants and study nurses, eligible participants were randomised (1:1) to either cognitive training and L-dopa administration or cognitive training and placebo. Age, sex, and the score on Raven’s Progressive Matrices were used as stratifiers. The randomisation was conducted using label shuffling with post-hoc non-parametric tests for the stratifiers and was run separately for each of the five waves using an R-script available at https://github.com/alex-lebedev (“RBTII” repository). All participants and all staff involved in administering the drug, cognitive training, and outcome assessments were masked to group assignment. To achieve masked drug administration, orange juice was mixed with L-dopa or administered as placebo. This was prepared and labeled by a nurse who was not involved in any other aspect of the study.

### Procedures

The study implemented an intervention period of four weeks, with five visits each week (~2.5 hours per visit, Monday-Friday), resulting in a maximum total of 20 intervention visits. All procedures were identical for the intervention group and the control group. Upon arrival, participants were given orange juice (with or without 100/25 mg of L-dopa/benserazide, trade name: Madopark® Quick mite. Roche, Basel, Switzerland) by a nurse who was masked to the group specification. After 45 minutes, during which time questionnaires evaluating mood, motivation, alertness and sleep were completed, participants commenced the cognitive training, which lasted for approximately 60 minutes. All participants then remained at the clinic for an additional 45 minutes for observation. The time window between the administration of the orange juice and the start of the cognitive training (45 min) was aligned with expected peaks in drug concentrations and effects. Similarly, total visit duration (2.5 hours) was motivated by drug elimination curves, according to which plasma concentrations of the drug are negligible 2.5 hours after the administration of L-dopa in combination with peripheral decarboxylase inhibitor (22). Dose selection was motivated by the results from two longitudinal studies completed in the healthy younger population that demonstrated positive effects of the drug on learning and good tolerability (7, 16). The cognitive training was designed according to the current recommendations in that it was adaptive in nature, targeted more than one construct (updating and switching), and promoted process-based as opposed to strategy-based improvements by including several training tasks and varying stimuli sets (23). The protocol incorporated three exercises: one focused on the ability to flexibly switch between different tasks (task-switching) and two others on the ability to continuously maintain and update mental representations (running span and n-back). Study outcomes included cognitive measures and Magnetic Resonance Imaging (MRI) data collected in five behavioural and one MRI session in the week before (pretest) and the week after the intervention period (posttest).

### Outcomes

The primary behavioural outcomes were: (1) spatial fluid intelligence, as a latent variable (see Statistical Analysis) measured by the Raven’s Progressive Matrices, the Wechsler Abbreviated Scale of Intelligence, and the BETA-III matrix reasoning test; and (2) verbal fluid intelligence, as a latent variable measured with the Analogies Task from Berlin Intelligence Structure Test, Syllogisms, and the Verbal Inference Test from the ETS Kit (See **Supplement S1** for complete list). Selection of the primary outcomes was motivated by previous literature suggesting a possibility that working memory training can produce improvements in measures of fluid intelligence (24).

Secondary cognitive outcomes were measures of working memory, episodic memory, and task switching ability. Structural brain imaging data were collected as a further secondary outcome. MRI scanning session (MRI Center, Huddinge Hospital) on a 3 Tesla scanner Siemens MAGNETOM Prisma equipped with a 24-channel research head coil was performed at the end of pretest and posttest weeks.

Six months later, participants were invited back to complete the cognitive assessment once more. Subjects were not unblinded until after the last follow-up visits.

Adverse events were assessed according to the most recent guidelines for Good Clinical Practice (European Medicines Agency, December 1^st^ 2016) and entailed comprehensive evaluation of their severity and possible connection with the drug. For detailed description, see study protocol at https://osf.io/nwwx8.

In order to quantify effective concentrations of the investigated drug, subjects’ blood samples were collected in the ethylenediaminetetraacetic acid-treated tubes at the first and last cognitive training visit, approximately 30-40 minutes after completing the training, to evaluate plasma levels of L-dopa and homovanillic acid. Plasma separation was performed within 4-5 hours via a 30-minute centrifugation at 3000×g. Plasma samples were stored in 1 ml aliquots at −80°C. The chemical analysis of L-dopa and homovanillic acid was performed with high-performance liquid chromatography analysis (Nexera-i HPLC system, Hertogenbosch, Netherlands; Antec electrochemical detection system, Leiden, Netherlands) and was blinded to the study groups.

### Statistical Analysis

Approximate power analysis was performed prior to the study launch with G*power 3, estimating the required sample size for detecting a group by time interaction (mixed ANOVA, F-test) on the primary outcomes, assuming a net standardised effect size (improvement for active group – improvement for control group) of 0.3 standard deviations, a test-retest stability coefficient of 0.70, and an alpha level (threshold for statistical significance) of 0.05. With these assumptions, we estimated a sample of 56 subjects as sufficient for detecting a true group × time interaction with a statistical power of 0.80. Considering a Bonferroni-corrected threshold for statistical significance (0.05/2 primary outcomes = 0.025), we aimed for a total of 64 subjects, which results, under the premises described above, in a statistical power of 0.786.

The main analysis used an implementation of structural equation modeling with latent change score modeling (25) to test the effect of L-dopa versus placebo on the outcomes of cognitive training (see **Supplement**, **Figure S2 and Table S3**). The analysis was implemented in the ‘lavaan’ package (26) within the R programming language environment, version 3.3.2 (2016-10-31). Latent variables were formed similarly for pre- and post-test assessments based on shared variance from multiple tests measuring each specific construct, and a latent change score, which represented the difference between the assessments, was estimated. Estimating intervention-related changes in this way, by forming a latent variable out of several tests, has the advantage of reducing the influence of measurement error and task-specific variance on the outcome measure, and hence also biases (e.g., regression to mean) that may affect raw change scores. The change factor was regressed on the group predictor (L-dopa/Placebo; dummy coded 1 vs. 0). The regression effect indicates the effect of experimental group on latent change from pretest to posttest (i.e., a time by group interaction). Prior to estimation, we z-standardised all variables, such that the size of this effect corresponds to the difference in gains over time between the two groups expressed in standard deviations. A separate model was estimated for each of the considered primary outcomes: spatial and verbal fluid intelligence.

Before model estimation, we cleaned and screened the data for outliers using the outlier labeling rule multiplying the interquartile range by a factor of 2.2. Detected outliers were deleted using pairwise deletion and the resulting missing values were accommodated under the missing-at-random assumption using full information maximum likelihood (FIML) estimation. Non-normally distributed variables were transformed employing applicable transforms until normality assumptions were met. We included all available data. Prior to hypothesis testing, measurement invariance assumptions were evaluated to ensure that the same latent variables are represented on each measurement occasion (27). Both models that incorporated the primary outcomes, spatial and verbal intelligence, met criteria for strict invariance. The same strategy was employed to analyse the follow-up data.

Training progress was analysed employing linear modelling that compared average level reached over the course of training (nested in tasks).

Plasma concentration of L-dopa and homovanillic acid (HVA) were analysed adhering to standard protocols of high-performance liquid chromatography (HPLC) and electrochemical detection (See **Supplement S4** for detailed description). Statistical analyses were conducted employing linear modelling (Group, Group × Visit effects on L-dopa/HVA levels, within-subject, random intercepts).

Structural MRI images underwent standardised steps for bias-field correction, segmentation, spatial normalisation and smoothing (FWHM of 8 mm) as implemented in the CAT12 (*http://www.neuro.uni-jena.de*), an SPM12 (*http://www.fil.ion.ucl.ac.uk*) toolbox installed in the MATLAB 2016 environment. See **Supplement S5** for more detailed description. Normalised and modulated grey matter probability maps were analysed employing mass-univariate within-subject ANOVA estimating group × time as a primary effect-of-interest. Yielded statistical parametric maps were adjusted for multiple tests employing a family-wise error-correction procedure. This was accomplished by testing the data against an empirical null distribution of maximum cluster size across 10,000 Gaussian noise simulations with an initial cluster-forming threshold of p<0.005. Clusters with expected false positive rate of <5% of (P_FWE_<0.05) were considered significant.

## RESULTS

Between January 1^st^ 2017 and October 10^th^ 2017, we screened 235 subjects, 64 of whom entered the study. Out of 64 recruited participants, 62 completed the study and MRI scans were collected for 57 of them (**Figure 1**). The scans were not obtained for five subjects because of psychological or physical discomfort that interfered with scanning sessions (neck problems, large head, claustrophobic reaction). Drop-outs were limited to two participants: one dropped-out on day 1 of the pre-test assessment (before randomisation), because he/she found the tasks too difficult, whereas the other participant, who was in the control group, dropped out on day 5 due to private commitments. **Table 1** summarises the background characteristics as a function of group. As expected, demographic data and baseline cognitive scores were similar in the two groups. Both groups completed equivalent number of training sessions (Placebo: 18.43 (16–21), L-dopa: 18.19 (14–21), two-sample Wilcoxon test, W = 500.5, p=0.77).

**Figure 1.**
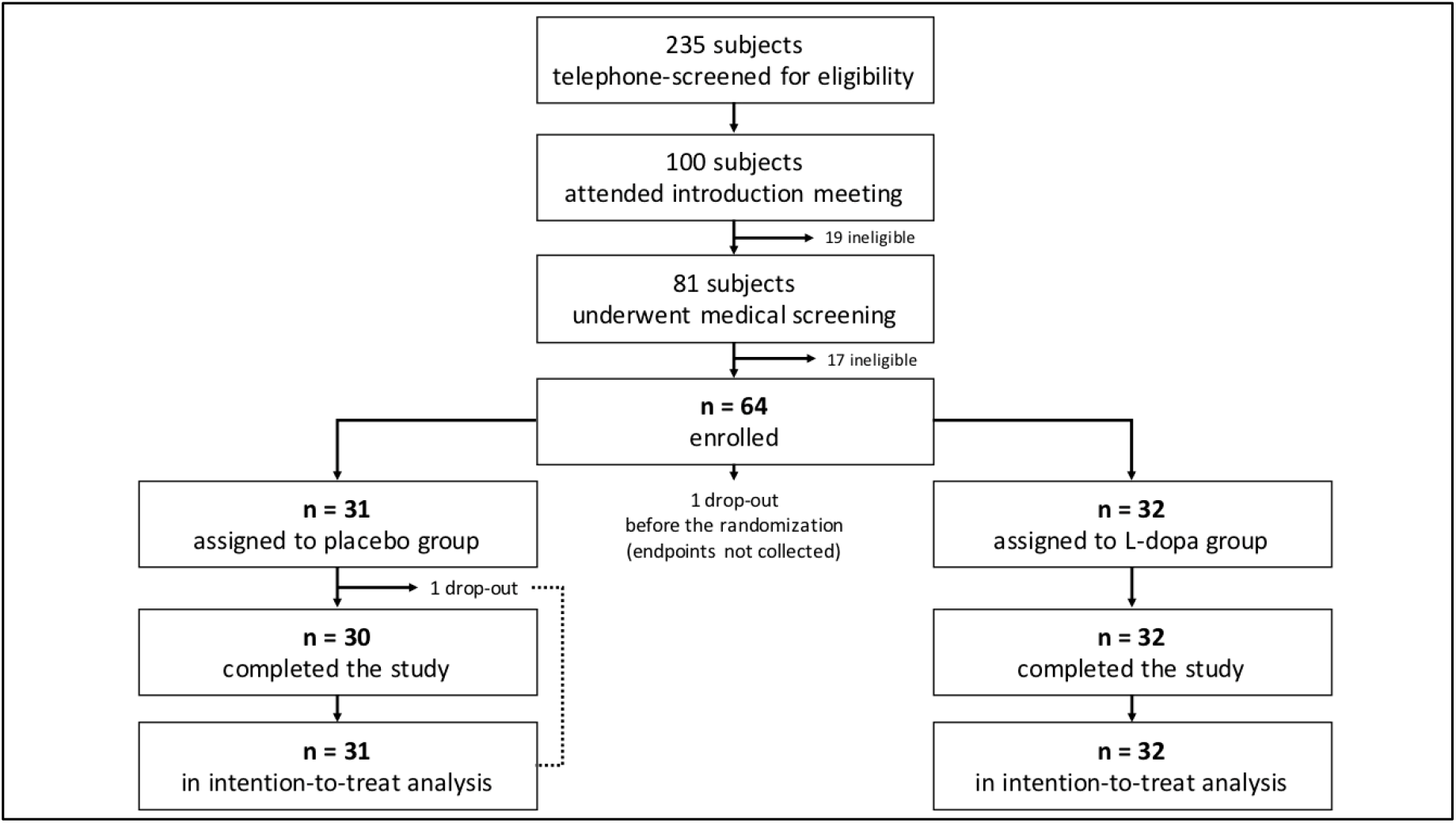
Trial profile. Out of 235 telephone-screened individuals, 64 entered the study. The primary outcomes were collected for 63 subjects: n=31 and n=32 in the placebo and L-dopa groups, correspondingly. A total sample of n=63 was used in the main analysis adhereing to intention-to-treat scheme.

**Table 1.** Baseline characteristics of the experimental groups.

|  | **Placebo**<br>(*n*=31) |  | **L-dopa**<br>(*n*=32) |  |
| --- | --- | --- | --- | --- |
|  | *Mean* | *SD* | *Mean* | *SD* |
| Age, years | 69.65 | 3.27 | 69.47 | 2.03 |
| Sex, f/m | 17/14 |  | 19/13 |  |
| MMSE score; median (range) | 30 (28-30) |  | 30 (26-30) |  |
| Education*; median (range) | 3 (0-10) |  | 3.5 (0-8.5) |  |
| BMI | 25.22 | 3.06 | 25.43 | 3.99 |
| SBP, mmHg | 139.87 | 11.65 | 137.13 | 12.05 |
| DBP, mmHg | 79.27 | 7.29 | 77.65 | 7.17 |
| Raven's Progressive Matrices score | 6.84 | 2.96 | 6.87 | 2.50 |
| Session, morning/afternoon | 16/15 |  | 18/14 |  |
| Drop-out, *n* subjects | 1 |  | 0 |  |
| MRI data collected, *n* (%) | 28 (90.32%) |  | 29 (90.63%) |  |
| 6-month follow-up available, *n* (%) | 24 (77.42%) |  | 27 (84.37%) |  |

\* - years after high-school;
*f/m* – female/male ratio; *BMI* – body-mass index; *SBP/DBP* – systolic/diastolic blood pressure; *MMSE* – Mini Mental State Examination.

Analyses of primary outcomes revealed that change of spatial fluid intelligence differed significantly between the groups, with the L-dopa group improving less compared to the placebo group between pre-test and post-test (Group × Time: standardised effect size −0.267 SDs, 95% CI [−0.498, −0.036]; p = 0.024; and **Figure 2**). Change of verbal fluid intelligence scores did not significantly differ between groups (Group × Time: standardised effect size, −0.081 SDs, 95% CI [−0.242, 0.080]; p = 0.323). Of note, traditional linear mixed analyses on unit-weighted composites of the primary outcomes showed essentially the same results as those we report here: spatial fluid intelligence, t(60) = 2.16, p = 0.03, and verbal fluid intelligence, t(60) = 0.11, p = 0.91.

Six-month follow-up data collected for a subset of 51 subjects revealed that the observed between-group differences in the spatial fluid intelligence improvements were still present 6 months after the intervention (standardised effect estimate: −0.371, 95% CI [−0.62, −0.122], p = 0.004). No statistically significant difference was found for verbal fluid intelligence.

No significant between-group difference was found for any of the secondary cognitive outcomes (See **Supplement Table S3**). Individual test scores (means and standard deviations) are available in the **Supplement Table S7**.

**Figure 2.**
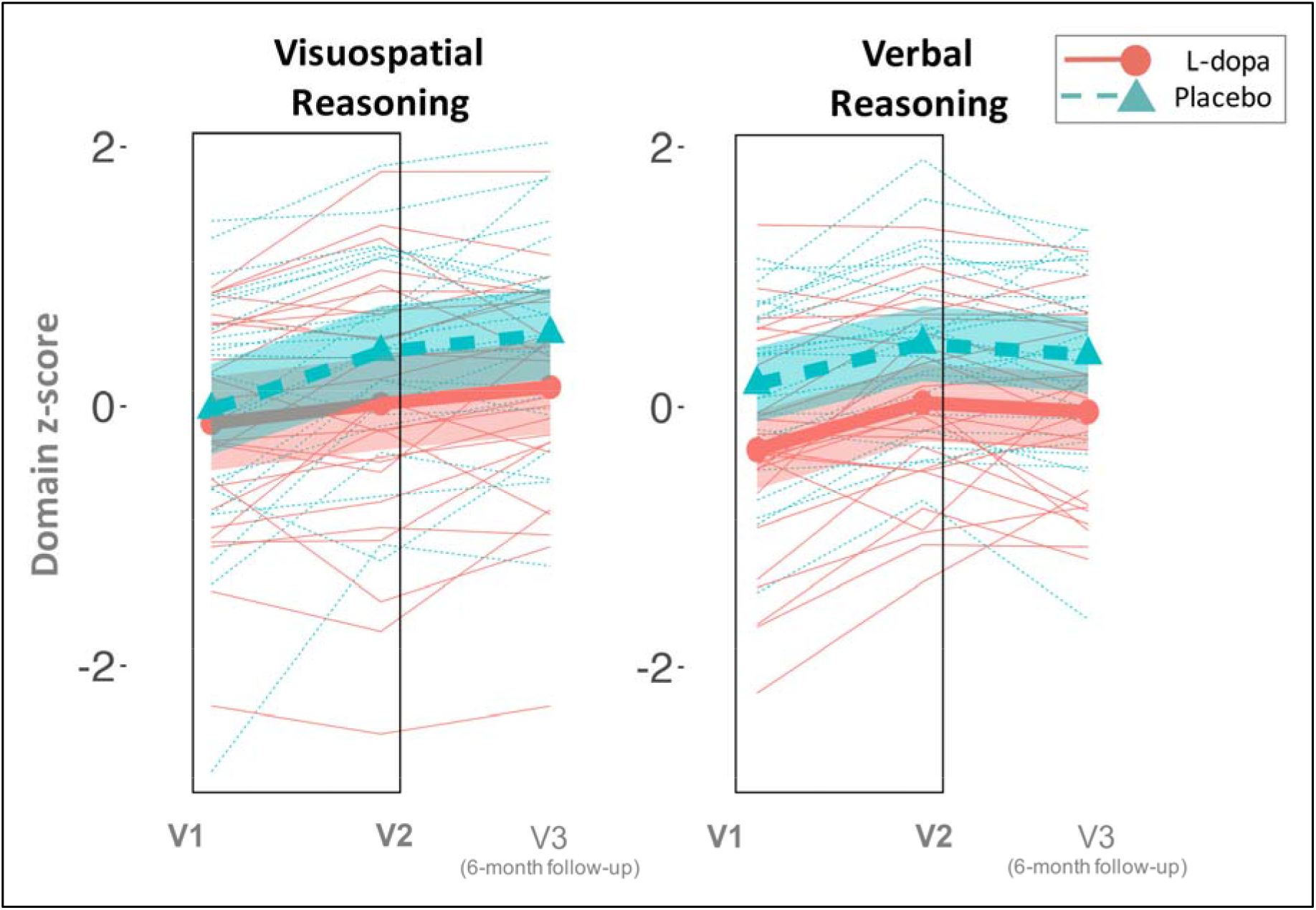
Pre-post changes in primary outcomes. Compared to the placebo group, subjects receiving L-dopa during a four-week working memory training program improved less in spatial reasoning domain. Shading represent 95 % CI. Boxes represent the preregistered timeline for the main analysis that intends to compare pre-post differences in performance. V1,2,3 – Visits 1,2,3

Between-group differences in training progress over the course of the intervention supported the main findings with the control group reaching higher difficulty levels across all three trained tasks (t(60) = 1.99, p = 0.05, **Figure 3**).

**Figure 3.**
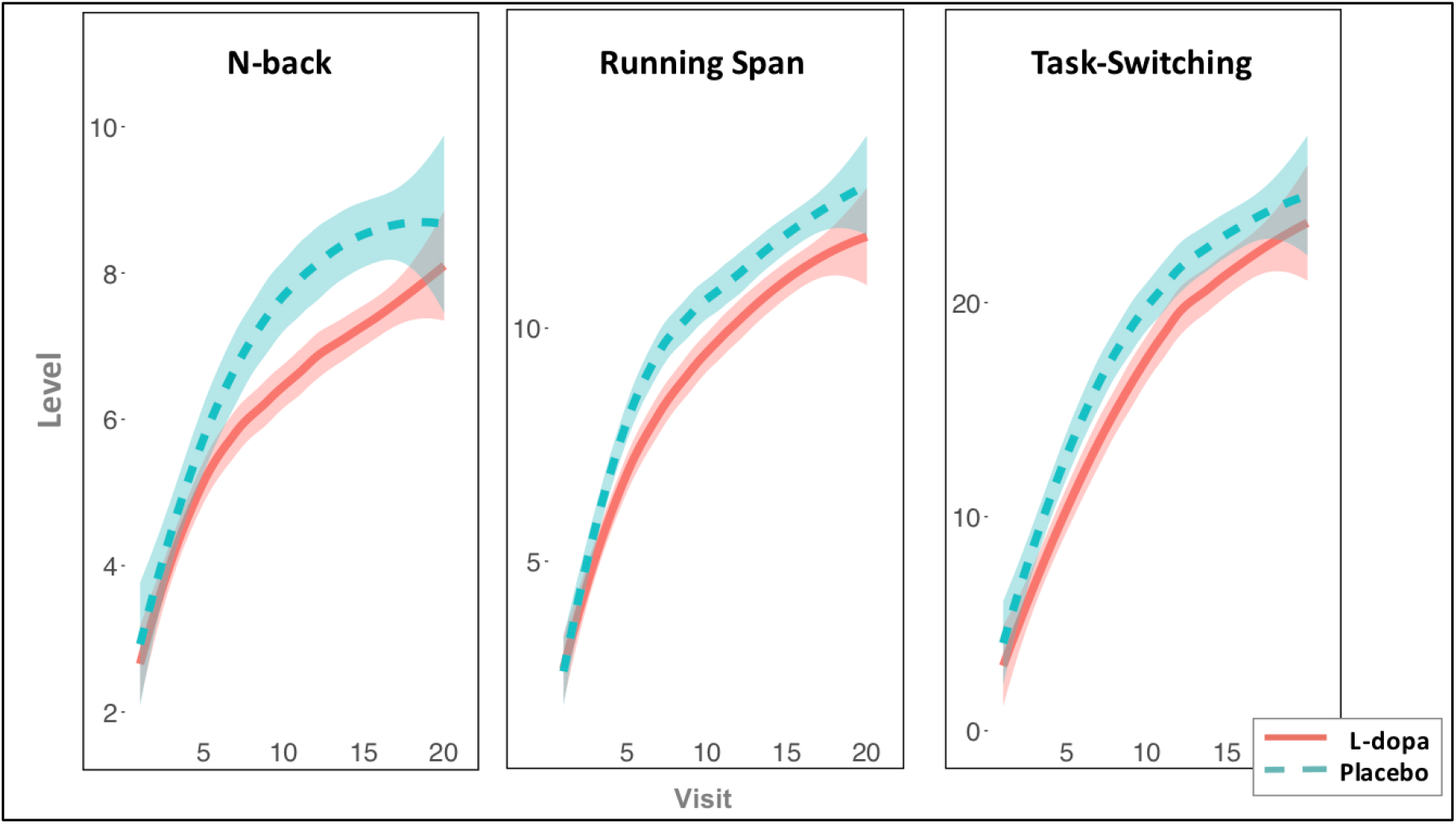
Training progress. X-axis: visit number; Y-axis: reached difficulty level Compared to the placebo group, subjects receiving L-dopa during a four-week working memory training program reached lower difficulty levels in all tasks. The lines are fitted with locally weighted scatterplot smoothing, shaded areas represent 95% CI.

Estimation of Group × Time effects on brain morphometry yielded a single significant cluster located in the midbrain (P_FWE_ < 0.05; MNI coordinates of the peak: −12 −24 −6 mm; see **Figure 4**). Matching it with the normalised high resolution delineations of the midbrain revealed an overlap with the substantia nigra (**Supplement S5**).

**Figure 4.**
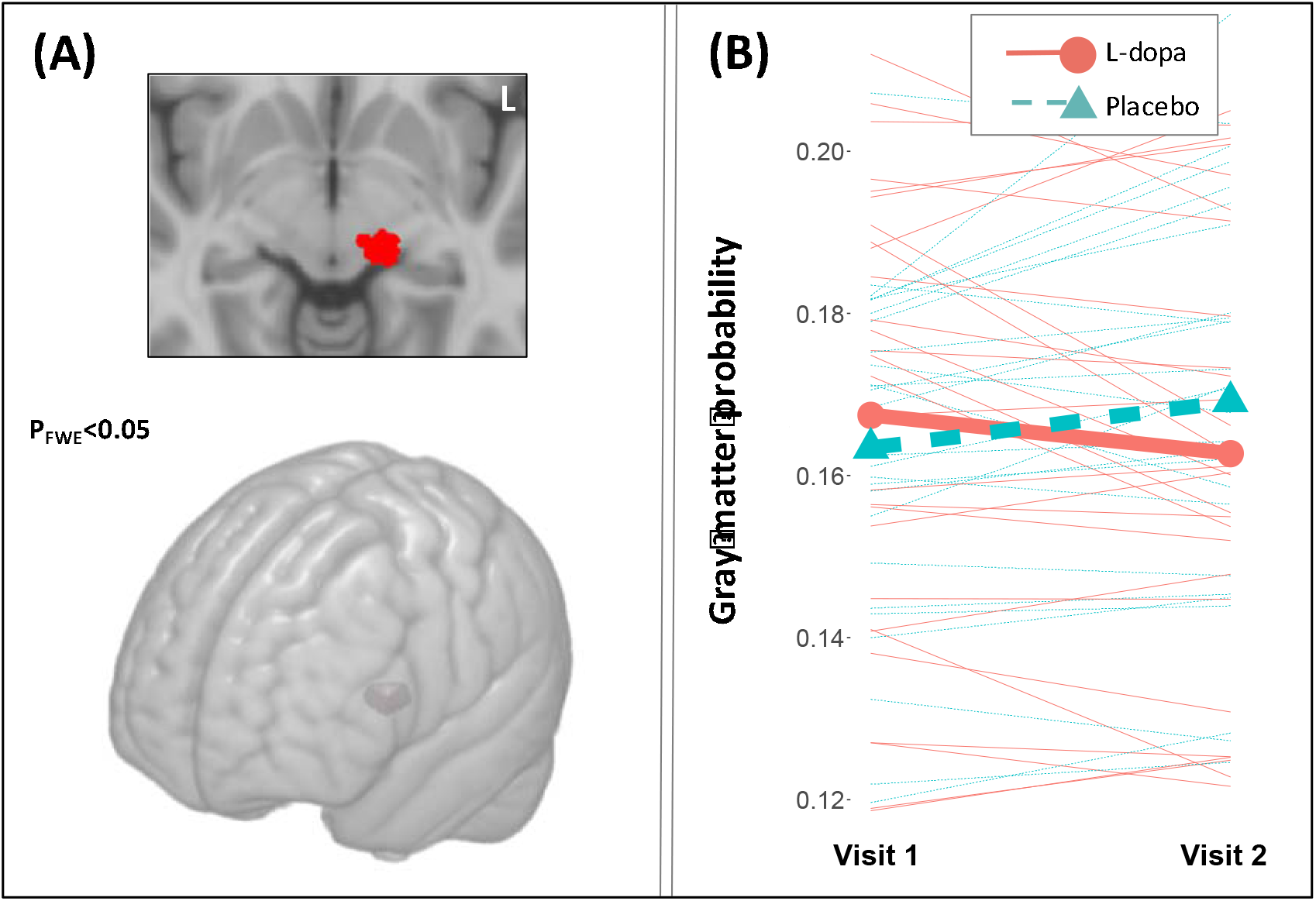
Structural Brain Changes. (A) Family-wise error-corrected (P_FWE_<0.05) cluster depicting Group × Time F-contrast; Statistically significant between-group differences in grey matter volumetric change observed in a midbrain region; (B) Between-group differences in volumetric changes.

When analysed separately, the structural changes in this region were significant in both groups. Specifically, control subjects showed increases in grey matter probability t(26) = 3.02, p = 0.006, whereas the L-dopa group exhibited reductions t(27) = −2.26, p = 0.032.

Concentrations of L-dopa and HVA were within the expected ranges (L-dopa: 2-1000 ng/ml; HVA: 1–500 ng/ml). As expected, Log_10_-transformed concentrations of L-dopa and homovanillic acid were higher in the L-dopa than in the placebo group: t(60)=15.01, p<0.001 and t(60)=9.96, p<0.001, correspondingly (See **Supplement S6** for more details). In addition, a significant 1.04 SDs increase in L-dopa concentrations was observed at the last training/intake day in the active group compared to the first administration, t(59)=15.11, p<0.001).

Furthermore, in the active group a negative relationship was found between plasma levels of the drug and improvements in visuospatial reasoning (i.e., those who had larger effective concentrations of L-dopa tended to improve less in the primary outcome of spatial reasoning; t(30)=2.06, p=0.048, **Figure 5**). It is also worth noting that the moderating effect of the body-mass index (BMI) on improvements in visuospatial reasoning was non-significant in the active group (Time × BMI interaction: t(29) = 0.25, p = 0.8) presenting no evidence for overdosing. It is also worth mentioning that BMI range in our sample was 19.8 - 37.9 without any extremely under-(BMI<16) or overweight (BMI>40) subjects.

**Figure 5.**
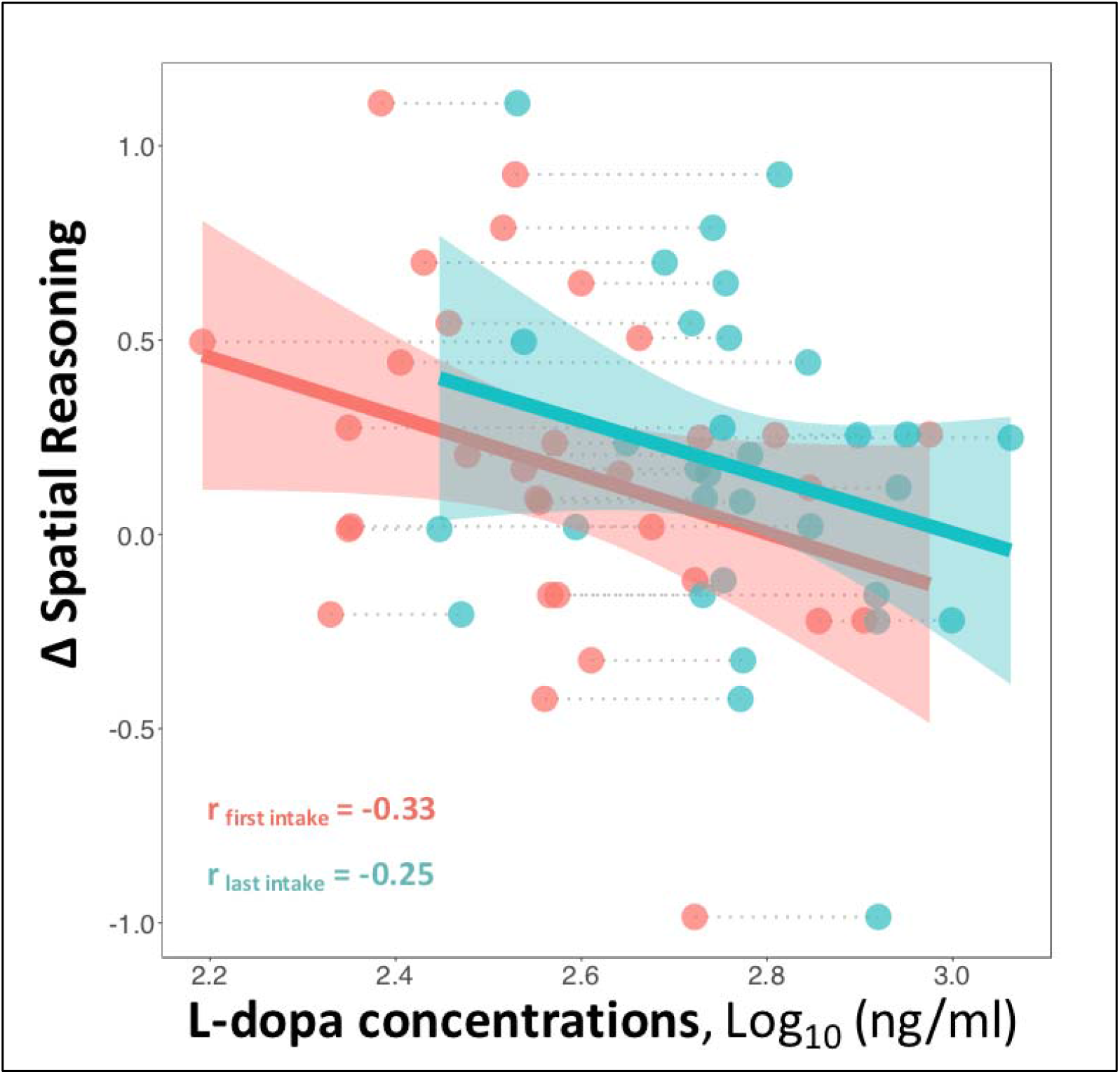
Correlation between plasma levels of L-dopa and changes in visuospatial reasoning in the active group. The plot shows that subjects who had larger plasma levels of the drug tended to improve less compared to those who had lower effective concentrations of L-dopa.

Thirty-one adverse events (AEs, 11 in the placebo and 21 in the L-dopa group) occurred in 17 subjects over the course of training. Typical AEs were common cold and related conditions (11), mild pain (7), mild vertigo/nausea (4). Total number of subjects with AEs did not significantly differ between the groups (Placebo: 7 subjects, L-dopa: 10 subjects, χ^2^ = 2.27, df = 4, p = 0.68); same was true for the drug-related AEs (Placebo: 1 subject, L-dopa: 2 subjects, χ^2^ = 0.98, df = 2, p = 0.61). Subjects’ quality of sleep was evaluated over the course of the study with the Karolinska Sleep Questionnaire and did not yield significant between-group differences for any of the subscales. Similarly, no significant between-group difference was found for mood and motivation over the course of training. Post-hoc evaluation of masking success revealed chance-level proportion of correct guesses in L-dopa (55%) and placebo (40%) groups.

## DISCUSSION

We did not find evidence for beneficial effects of exogenous augmentation of dopaminergic neurotransmission on cognitive performance and learning in healthy older adults. On the contrary, subjects receiving L-dopa improved less on visuospatial fluid intelligence, a primary outcome of the training intervention, after four weeks of working memory training compared to those who received placebo treatment. Subjects receiving L-dopa also progressed worse during training when compared to placebo subjects. The observed between-group differences in visuospatial fluid intelligence were still statistically significant 6 months after the intervention. The groups also demonstrated opposite direction of structural changes in a midbrain region, overlapping with the template location of substantia nigra, which is a key regulatory region rich in dopamine neurons involved in learning and plasticity (28, 29). Specifically, the control group exhibited increases of grey matter volume in this region whereas the group that received L-dopa showed reductions.

Although post-hoc, we interpret our results to suggest that the exogenous administration of the dopamine precursor may have perturbed a balanced dopaminergic system that previous studies has shown to be involved in training-related effects on performance (15). This was also supported by negative correlation between training-related improvements in performance and plasma concentrations of the drug observed in the active group. The differential volumetric changes in the dopamine-rich region further support this interpretation. Thus, whilst our results are not at odds with basic science that clearly implicates dopamine in learning and plasticity (10, 12, 13), they indicate that exogenous administration the dopamine precursor is rather negative than beneficial for the dopaminergic system in the healthy aging brain. We also note that, despite some positive results from repeated L-dopa administrations on verbal learning domain reported in previous studies of younger adults (7, 16), our negative findings are in line with a recent study that showed detrimental effects of the same L-dopa dose on reward reversal learning in younger and older healthy adults, which were of the same magnitude irrespective of initial cognitive performance and expected baseline dopamine levels (30). Further studies are clearly needed to clarify under what circumstances and for which cognitive domains pro-dopaminergic pharmacological interventions may have differential effects, especially in the light of recent findings showing clear regional heterogeneity of age-related decline in dopamine receptor availability (31).

More generally, the negative response to pro-dopaminergic medication observed in the present study suggests that cognitive decline in healthy aging is a complex and multifactorial process, a process that appears to result in balanced physiological states, which, whilst being associated with reduced overall self-regulatory capacity (32), still preserve mechanisms of homeostatic plasticity (33–35), which implies that activity within the dopaminergic circuit is regulated by intrinsic feedback loops to maintain optimally balanced levels of the neurotransmitter thereby preventing excessive increases in excitatory activity. The negative effect observed in the present study appears to be the opposite of what is known as “dopamine supersensitivity” typically present in dopamine-deficient conditions (36–38). Translating this finding further to clinical and cognitive neuroscience, it is worth noting that although the dopamine system is affected in several ways in aging (15, 18, 31, 39), acute pro-dopaminergic supplementation appears to perturb a balanced system, with negative behavioural consequences. This is fully in line with the results from positron emission tomography studies showing that despite apparent negative effects of age on dopamine transporters and receptor density in the healthy individuals, its synthesis capacity remains relatively unaffected (39). It is also consistent with recent evidence indicating that the balance between dopamine release and receptor density is critical for cognitive performance (40). In age-related neurodegenerative diseases, on the other hand, disruption of the homeostasis has reliably been observed and may even be the core pathophysiological abnormality triggering development and deterioration of cognitive functions (41). Thus, it appears clear that aging should not be approached as a disease per se, but rather as a physiological process associated with a gradual decline in many interconnected biological and behavioural capacities. Maintenance of the aging organism may therefore be best achieved by early investments in healthy lifestyle and multi-modal interventions affecting it (1, 2).

Our results warrant more studies about the effects of L-dopa on brain and cognition in clinical populations, which often involve substantially higher dosages and longer time periods. Indeed, even though dopamine replacement therapy has been shown to be successful for counteracting motor impairment associated with Parkinson’s disease, studies that explore its effects on cognition have yielded mixed results (17, 42). The negative effects of L-dopa supplementation on learning and midbrain structures observed in our study present an urgent need to carefully investigate the longitudinal course of brain changes in de-novo Parkinson’s patients, for which early dopamine replacement therapy is currently a subject of debates (43, 44).

Some important study limitations need to be addressed. The most important one is the absence of additional control groups not receiving any cognitive training interventions. This limits interpretation of the results, as with only two groups we cannot unambiguously infer that the between-group differences are driven by the interaction of L-dopa with cognitive training and not by direct and prolonged effects of the drug on cognitive abilities or an interaction with re-test effects (which would still entail effects on learning but of a different kind). However, elimination of L-dopa is fast (blood concentrations are expected to be negligible when the subjects leave the training facility), and all subjects had minimum 24-hour washout period and were medication-free on pre- and post-testing days. This, however, does not completely rule out a possibility of a cumulative effect on the brain concentrations, which, in turn, may drive the observed detrimental effects on performance. In this context, it is also worth mentioning that L-dopa concentrations were significantly higher in the active group at the last compared to the first intake. We interpret this finding as due to accelerated gut absorption of the drug previously reported in animal studies with repeated L-dopa administration (45). Nevertheless, similar to the main results, the placebo group reached higher difficulty levels over the course of training in all tasks. We think that these learning-related effects are the most likely explanation of the performance differences at post test, which may also explain why they were still present at long-term follow-up conducted 6 months after the intervention (at a point when differential L-dopa concentration between the groups are unlikely).

It is also important to acknowledge limitations of the MRI-derived measures of grey matter probability. The differential changes in the midbrain were observed in measures that are derived from T1-weighted images, which are known to be highly sensitive to pharmacological manipulations (46). We cannot determine whether the observed changes are due to true volumetric changes or whether they are a result of relatively transient changes in for example blood flow. It is worth noting that the increases observed for the control group in the present study are consistent with previous reports of changes in the dopaminergic system induced by cognitive training (10). In line with this background, we also hypothesise that the observed effects in the L-dopa group may reflect reactive changes in the dopamine system in response to repeated administration of an exogenous precursor of the neurotransmitter.

In addition, even though our literature review indicated that the selected dosage was appropriate to induce the effects of interest with minimal side-effects, we cannot completely rule out the possibility that other drug amounts may lead to different results. However, a negative and linear correlation between plasma concentrations of the drug and improvements in visuospatial reasoning observed in the active group, as well as absence of any moderating effects of the body-mass index on the aforementioned improvements make this possibility unlikely. Finally, despite the fact that our study is well-powered and the largest of its kind (7, 16) an additional caution must be advised for making direct generalisations to broader populations, especially to patient groups discussed above.

We conclude that daily L-dopa supplementation does not enhance cognitive performance and learning during cognitive training in healthy older adults and may in fact have disadvantageous effects. Our findings raise serious concerns about usefulness of novel L-dopa-containing supplements that claim to have neuroprotective and learning-enhancing properties and suggest that caution is needed with regard to early dopamine replacement treatment interventions in neurological disorders, encouraging more rigorous evaluation of their effects on the brain and cognition in populations that often receive the drug in larger doses over long periods.

## CONTRIBUTORS

ML conceived the study. ML, AL, JN and MK designed the trial. AL, ML, and MK coordinated the trial. AL and ML analysed the behavioural and MRI data. DB and EL analysed the blood samples. AL, ML, JN interpreted the results. GS, PA, JL, WF and CH collected the data. ML obtained funding. All authors revised the paper for important intellectual content.

## ACKNOWLEDGMENTS

First and foremost, we thank all participants of the REBOOT-II project. We also thank Marie Helsing for help with administrative work, recruitment and organisation. Thanks also to Aleksandra Lebedeva, Erika Bereczki, Marc Guitart-Masip, Lars Bäckman, Patrik Fazio, and Yvonne Brehmer for the valuable discussions and comments on the study plan and results.

## DATA AVAILABILITY STATEMENT

All of the analysis steps are documented in R and MATLAB scripts at https://github.com/alex-lebedev (“RBTII” repository). This trial was preregistered at the European Clinical Trial Registry, EudraCT # 2016-000891-54, and The Open Science Framework Registry, DOI 10.17605/OSF.IO/AAM9U. According to Swedish law the whole dataset and biological materials cannot be freely accessible, but can be requested from the authors for specific research projects. This requires a data transfer agreement, which effectively transfers the confidentiality obligations of the institution at which the original research was conducted to the institution of the recipient of the data.

