## Supplement for "Effects of daily L-dopa administration on learning and brain structure in older adults undergoing cognitive training: a randomised clinical trial"

***S1. Investigated cognitive domains and corresponding tasks***

| Domain | Tasks |
|---|---|
| **Spatial Intelligence** PE | Ravens progressive matrices, WASI-II matrices, BETA-III matrices |
| **Verbal Intelligence** PE | ETS kit verbal inference, BIS analogies, Syllogisms |
| **Updating** SE | N-back, Running span (untrained stimuli) |
| **Rule-Switching** SE | Spatial switching, Numerical switching |
| **Task-Switching** SE | Task-switching (3 difficulty levels, untrained stimuli) |
| **Updating-trained** SE | N-back, Running span (trained stimuli) |
| **Task-Switching-trained** SE | Task-switching (3 difficulty levels, trained stimuli) |
| **Episodic Memory** SE | Spatial recall, verbal recall |

PE = Primary endpoint; SE = Secondary endpoints

***Figure S2. Structural Equation Model***

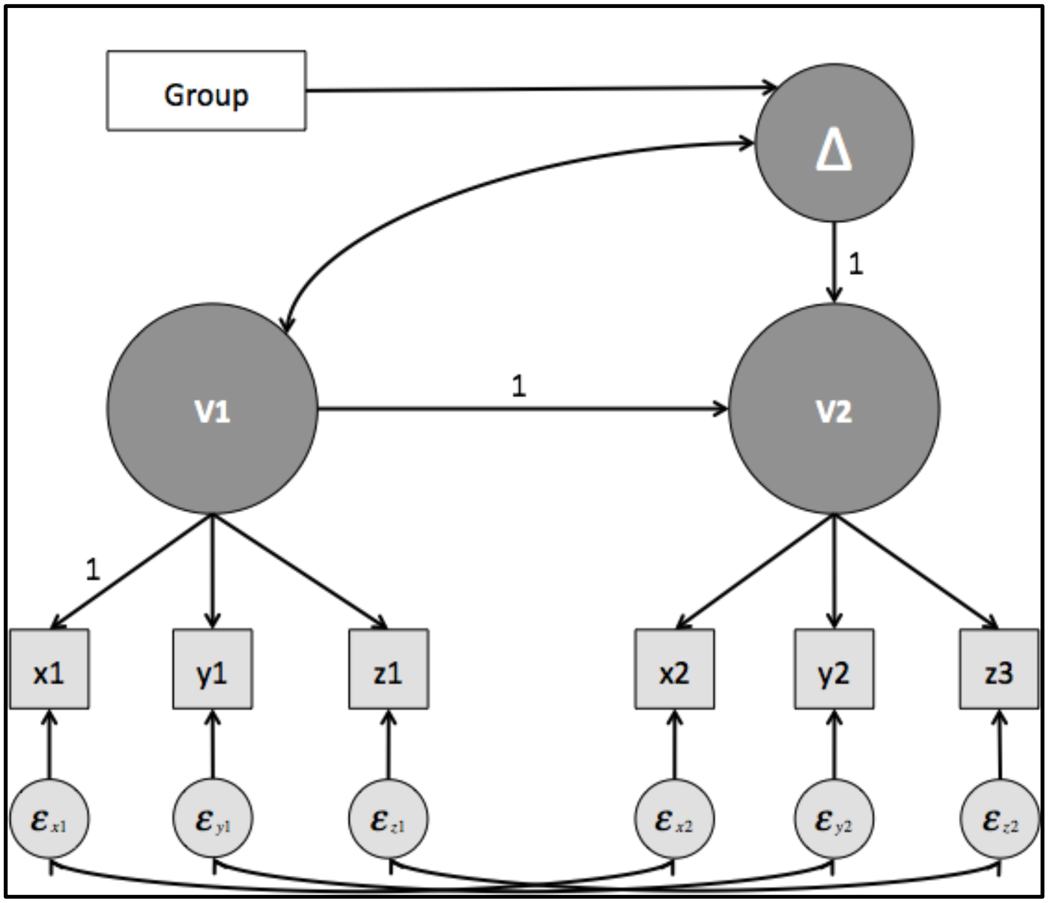


*A simplified illustration of the structural equation model used in the main analyses.*
*V1/V2 – Visits 1 (pre-test) and 2 (post-test); Δ – Latent change in cognitive performance;*
*ε – residuals; Box "Group" represents a factor that determines group affiliation (Placebo/L-dopa)*
*Measured variables (indicators x, y and z) are represented in squares (inward one-sided arrows represent loadings), whereas latent variables are shown in dark circles. Single-headed arrows represent regression (causal relationship), double-headed arrows represent non-causal relationships (equivalent to correlation).*

*Table S3. Between-group differences in all cognitive outcomes*

| Domain | Model | | Group effect (G~Δ) |
|---|---|---|---|
| | IA met | CFI/RMSEA | Estimate (p) |
| **Spatial Intelligence** $_{PE}$ | Strict | 0.97/0.083 | -0.267 (0.024)* |
| **Verbal Intelligence** $_{PE}$ | Strict | 0.957/0.079 | -0.081 (0.323) |
| **Updating** $_{SE}$ | Strict | 0.982/0.058 | -0.048 (0.707) |
| **Rule-Switching** $_{SE}$ | Strict | 0.946/0.116 | -0.148 (0.265) |
| **Episodic Memory** $_{SE}$ | Strict | 0.953/0.110 | -0.173 (0.278) |
| **Task-Switching** $_{SE}$ | Weak | 0.99/<0.001 | 0.315 (0.09) |
| **Updating-trained** $_{SE}$ | Strong | 0.828/0.232 | -0.116 (0.249) |
| **Task-Switching-trained** $_{SE}$ | Weak | 0.99/0.023 | -0.217 (0.127) |

PE = Primary endpoint; SE = Secondary endpoints; IA – Invariance Assumption

### *S4. Detection of plasma levels of L-dopa and homovanillic acid*

Plasma concentration of L-dopa and homovanillic acid (HVA) were analysed employing high-performance liquid chromatography (HPLC) and electrochemical detection according to standardised procedures. All chemicals were of analytical grade unless otherwise stated. The lab water was derived from an ELGA Purelab Flex water purification system from Veolia Nederland (Nieuwegein, the Netherlands). Methanol (ULC-MS grade) was from Biosolve (Valkenswaard, the Netherlands). L-dopa, HVA, acetic acid, L-cysteine, sodium monohydrogen phosphate, sodium dihydrogen phosphate and octane-sulphonic acid were from Sigma-Aldrich (Zwijndrecht, the Netherlands). Phosphoric acid, ethylenediaminetetraacetic acid (EDTA) and ascorbic acid were derived from J.T. Baker (Deventer, the Netherlands).

L-dopa and homovanillic acid were dissolved and diluted in antioxidant solution. This solution contained 100 mM acetic acid with 3.3 mM L-cysteine and 0.3 mM EDTA. Before use of the antioxidant solution, ascorbic acid was freshly added in a concentration of 12.5 µM. L-dopa concentrations were from 2 – 1000 ng/ml and HVA concentrations were in the range 1 – 500 ng/ml. To 50 µl of blank plasma sample, 50 µl of calibration solution was added and vortexed briefly.

Plasma samples in a volume of 50 µl were mixed 1:1 with antioxidant solution. After mixing with 50 µl of 10 % trichloroacetic acid during 2 minutes, centrifugation was performed at 4,500 xg (10 minutes at 4 degrees Celsius). After mixing of 80 µl of supernatant with 40 µl of 1M Phosphate buffer (pH 5.5), centrifugation was performed at 20,000 xg for 10 minutes at 4 degrees Celsius. 80 µl of sample was inserted into vials and 25 µl was injected into the HPLC system.

The HPLC system consisted of a Nexera-i system from Shimadzu (Hertogenbosch, the Netherlands) which was coupled to an electrochemical detection system from Antec (Leiden, The Netherlands). The mobile phase consisted of ULC-MS grade methanol and a solution of 50 mM phosphate, 1.5 mM octane sulfonic acid and 0.3 mM EDTA at pH 2.9 in a ratio of 5.55 (v/v). An Inertsil column (250 x 2.0 mm) (Inacom, Overberg, the Netherlands) was operated at 0.2 ml /min and connected to a VT-03 electrochemical cell at 0.85V. The cell was equipped with a 25 µm spacer and operated at 30 degrees Celsius. The LC vials were kept at 8 degrees Celsius pending analysis.

***S5. Magnetic resonance imaging***

*Quality Control*

A quality control procedure was conducted blindly to study group information and consisted of three stages: 1) manual screening for artefacts and overall grey/white matter contrast quality, 2) an automated evaluation of the weighted segmentation quality, which takes into account image resolution, signal-to-noise ratio and magnitude of a field bias (weighted overall quality criterion had to be at least 80%, indicating "good" segmentation quality), 3) calculation of the sum of the squared distance of each normalised and smoothed grey matter image from the sample mean (the sum of the squared distance of an individual grey matter map had to be within 2 SDs from the sample mean). Two T1-weighted images (one from the placebo and one from the L-dopa group) did not pass the above-described quality control procedure at stages 2 and 3:

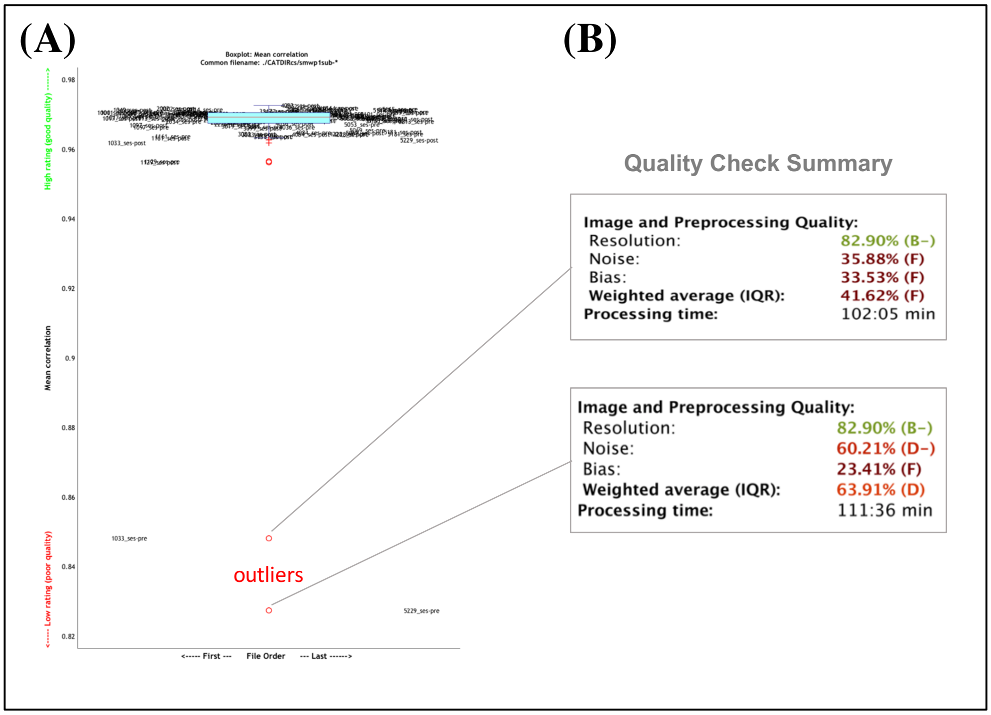


*Automated outlier detection procedure of the brain imaging data based on sample homogeneity detected two data-points outside the main sample cloud. Those were the same scans that had the worst preprocessing quality scores. They were excluded from the main analysis.*

*Scanning Procedures*

T1 3D brain scans were collected for a subsample of 57 subjects at one site (MRI Center, Huddinge Hospital) on 3 Tesla scanner Siemens MAGNETOM Prisma with a 24-channel research head coil. The procedure employed a standardised GRAPPA MPRAGE acquisition protocol according to Alzheimer's Disease Neuroimaging Initiative standards (ADNI-3). TR/TE = 2300/2.95 ms, Base resolution = 256, FoV read = 270 mm, Voxel size = 1.1 x 1.1 x 1.3 mm.

*Image preprocessing*

Structural MRI data processing was conducted employing standard voxel-based morphometry stream implemented in the Computational Anatomy Toolbox (CAT12; http://dbm.neuro.uni-jena.de/cat) based on the Statistical Parametric Mapping (SPM12; http://www.fil.ion.ucl.ac.uk/spm) software installed in the MATLAB 2016 environment.

All 3D T1-weighted MRI scans were normalised using an affine followed by a non-linear registration, corrected for bias field inhomogeneity, and segmented to extract a grey matter component [1]. Normalisation of grey matter density maps to MNI space was done using geodesic shooting [2]. In order to provide a possibility to compare the absolute amounts of tissue corrected for individual differences in brain size, a non-linear deformation on the normalised segmented images was performed yielding modulated tissue probability maps. The modulated and normalised maps were then smoothed with a full width at half maximum (FWHM) kernel of 8 mm. Resampled voxel size was 1.5 × 1.5 × 1.5 mm3. The implemented script can be accessed at https://github.com/alex-lebedev.

The resulting smoothed maps were analysed employing mass-univariate within-subject ANOVA estimating group × time as a primary effect-of-interest. Yielded statistical parametric maps were adjusted for multiple tests employing a family-wise error-correction procedure. This was accomplished by testing the data against an empirical null distribution of maximum cluster size across 10,000 Gaussian noise simulations with an initial cluster-forming threshold of $p<0.005$. Clusters with expected false positive rate of <5% of ($P_{FWE}<0.05$) were considered significant.

*Cluster Localization*

The resulting cluster of difference was matched with high resolution segmentation of the subthalamic area [3] accessed at Neurovault (https://neurovault.org/collections/550/).

We found that the cluster-of-difference partially overlapped with the substantia nigra:

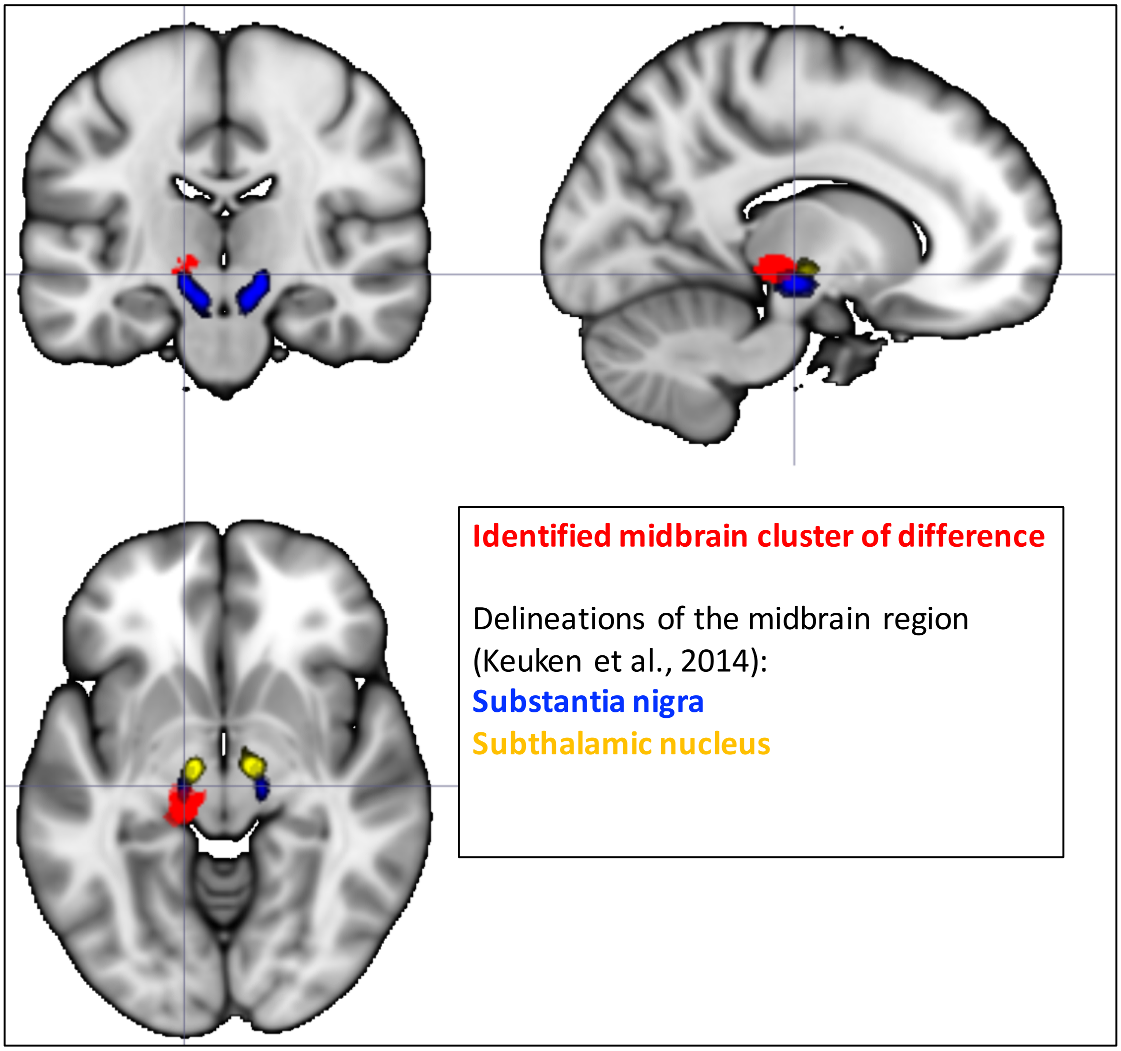


***S6. Analysis of L-dopa and homovanillic acid levels in plasma***

Between-group comparison of L-dopa and homovanillic acid levels in plasma revealed significant differences:

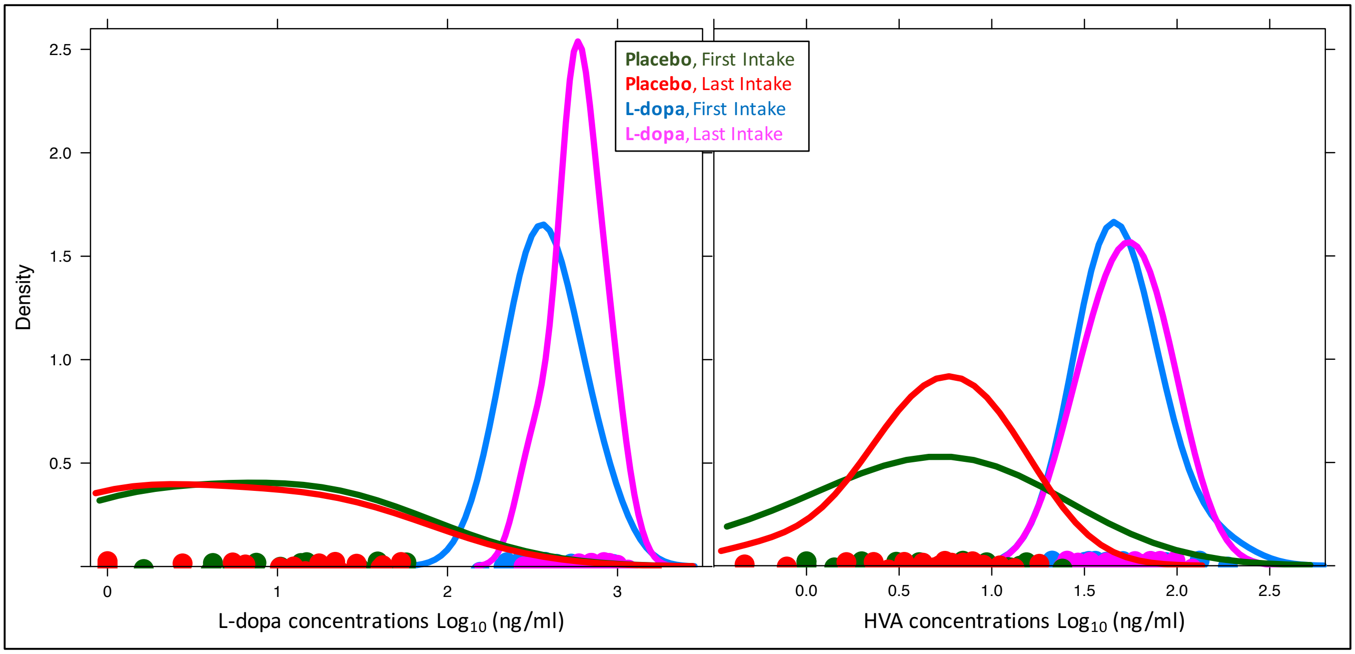


***Figure S5****. Plasma concentrations of L-dopa and homovanillic acid (HVA).*

*L-dopa*

Placebo: $Log_{10}$, ng/ml of, control group, visits 1/2 = 0.77±0.65 / 0.66±0.66

Treatment: $Log_{10}$, ng/ml of, control group, visits 1/2 = 2.57±0.18 / 2.76±0.14

Between-group difference: T(60) = 15.01, $p<0.001$

*Homovanillic acid*

Placebo: $Log_{10}$, ng/ml of, control group, visits 1/2 = 0.52±0.64 / 0.69±0.36

Treatment: $Log_{10}$, ng/ml of, control group, visits 1/2 = 1.70±0.19 / 1.72±0.18

Between-group difference: T(60) = 9.96, $p<0.001$

A significant 1.04 SDs increase in L-dopa concentrations was observed in the active group at the last compared to the first day of administration (Group x Visit effects on L-dopa levels, within-subject, random intercepts: T(59)=15.11, $p<0.001$).

**Table S7. Individual test scores (means and standard deviations)**

| TASK | VISIT 1 | | VISIT 2 | |
|---|---|---|---|---|
| | PLACEBO | L-DOPA | PLACEBO | L-DOPA |
| RAVENs | 6.84 ± 2.4 | 6.88 ± 2.5 | 8.43 ± 2.11 | 7.5 ± 2.69 |
| BETA | 17.42 ± 3.31 | 17.34 ± 3.33 | 18.2 ± 3 | 18.16 ± 3.52 |
| WASI | 21 ± 3.1 | 21.06 ± 2.47 | 21.9 ± 2.76 | 21.09 ± 3.32 |
| Verbal Inference | 11.35 ± 3.44 | 10.19 ± 3.22 | 13.03 ± 3.27 | 11.47 ± 3.12 |
| Word Comprehension | 24.48 ± 1.75 | 23.69 ± 2.28 | 24.53 ± 2.06 | 24.31 ± 2.19 |
| Syllogisms | 18.71 ± 3.73 | 16.56 ± 4.26 | 19.63 ± 3.3 | 17.94 ± 3.12 |
| Analogies | 6 ± 2.11 | 4.75 ± 1.98 | 6.47 ± 2.43 | 5.19 ± 2.32 |
| Near Updating (Level 2) | 1.86 ± 0.64 | 1.67 ± 0.51 | 2.37 ± 0.47 | 2.44 ± 0.37 |
| Near Updating (Level 4) | 2.68 ± 0.95 | 2.41 ± 0.92 | 3.49 ± 0.48 | 3.15 ± 0.78 |
| Trained Updating (Level 2) | 2.18 ± 0.57 | 2.22 ± 0.35 | 2.63 ± 0.33 | 2.67 ± 0.22 |
| Trained Updating (Level 4) | 2.97 ± 0.78 | 2.83 ± 0.72 | 3.64 ± 0.28 | 3.41 ± 0.5 |
| Trained Spatial Updating | 1.24 ± 0.48 | 1.03 ± 0.42 | 1.34 ± 0.59 | 1.31 ± 0.58 |
| Near Rule-Swtiching1 (cost) | 242.57 ± 168.24 | 201.64 ± 136.08 | 170.72 ± 107.63 | 148.23 ± 76.65 |
| Near Rule-Swtiching2 (cost) | 447.47 ± 218.16 | 399.66 ± 181.57 | 450.95 ± 220.62 | 468.6 ± 157.53 |
| Near Task-Switching, level 1 (cost) | 771.71 ± 257.28 | 812.99 ± 270.45 | 728.5 ± 170.39 | 824.69 ± 244.95 |
| Near Task-Switching, level 3 (cost) | 877.96 ± 232.36 | 928.82 ± 261.35 | 778.01 ± 182.79 | 875.88 ± 248.18 |
| Near Task-Switching, level 4 (cost) | 842.33 ± 206.48 | 881.03 ± 283.27 | 748.32 ± 147.6 | 843.19 ± 249.91 |
| Trained Task-Switching, level 1 (cost) | 943.43 ± 263.87 | 930.39 ± 278.6 | 736.53 ± 207.31 | 800.9 ± 211.28 |
| Trained Task-Switching, level 3 (cost) | 888.46 ± 268.37 | 882.95 ± 279.55 | 699.19 ± 199.34 | 766.91 ± 218.4 |
| Trained Task-Switching, level 4 (cost) | 727.47 ± 226.23 | 755.82 ± 261.61 | 670.66 ± 150.37 | 714.16 ± 215.73 |
| Numerical Flanker (cost) | 424.86 ± 184.88 | 399.14 ± 141.67 | 360.17 ± 137.66 | 341.7 ± 114.96 |
| Spatial Flanker (cost) | 359.96 ± 125.73 | 336.24 ± 157.77 | 316.36 ± 129.95 | 297.21 ± 136.72 |
| Verbal Recall | 15.67 ± 4.77 | 16.16 ± 4.14 | 18.1 ± 5.28 | 16.69 ± 5.6 |
| Spatial Recall | 13.23 ± 3.68 | 11.44 ± 3.58 | 15.3 ± 4.49 | 13.62 ± 2.98 |
| Near 2-back | 0.8 ± 0.12 | 0.79 ± 0.12 | 0.86 ± 0.13 | 0.87 ± 0.08 |
| Near 3-back | 0.68 ± 0.08 | 0.69 ± 0.08 | 0.73 ± 0.1 | 0.74 ± 0.06 |
| Trained 2-back | 0.79 ± 0.13 | 0.78 ± 0.12 | 0.87 ± 0.12 | 0.86 ± 0.1 |
| Trained 3-back | 0.66 ± 0.07 | 0.65 ± 0.06 | 0.74 ± 0.1 | 0.72 ± 0.09 |
